# Droplet microfluidics reports digital and tuneable single-platelet ADP secretion describing primed, active and hyper-active states

**DOI:** 10.64898/2026.07.24.740520

**Authors:** Max Saito, Theo Hornsey, Ahmed Elserwey, Simon Lane, Craig E. Hughes, Chris Jones, Nick Curzen, Jonathan West

**Affiliations:** Cancer Sciences, Faculty of Medicine, University of Southampton, UK; School of Biological Sciences, Faculty of Environmental and Life Sciences, University of Southampton, UK; Faculty of Medicine, University Hospital Southampton NHS Foundation Trust, UK; School of Chemistry, Faculty of Engineering and Physical Sciences, University of Southampton, UK; Institute for Life Sciences, University of Southampton, UK; School of Biological Sciences, University of Reading, UK; Coronary C Structural Heart Research Group, University Hospital Southampton NHS Foundation Trust

## Abstract

New platelet function assays that provide an accurate assessment of platelet reactivity as a surrogate for thrombosis risk could change the paradigm for acute myocardial infarction and stroke prevention. Considering ADP secretion as a hallmark of platelet activation, we developed a method for encapsulating single platelets with a fluorescent reporter for extracellular ADP in picolitre droplets. The method, termed the Droplet ADP Secretion Assay (DASA), robustly detected dose-dependent ADP secretion responses to agonists. Digital ADP secretion heterogeneity was observed with the major fraction of platelets responding and the remaining fraction being non-responders. Primed platelets had elevated responses to agonists, with increased ADP secretion and an increased fraction of secreting platelets, demonstrating that priming can convert otherwise inactive platelets to a reactive state. Uniquely, the method detected ADP secretion from primed platelets in the absence of agonists. In contrast, primed platelets were not detected by flow cytometry involving CD63 and CD62P endpoints. In a preliminary clinical study, some patients had platelets with a priming-like ADP secretion signature, suggesting treatment with P2Y_12_ inhibitors may be warranted. *In vitro* aspirin and ticagrelor treatments reduced ADP secretion responses to agonists. In summary, DASA offers a novel approach to assess platelet reactivity and holds promise for assessing thrombotic tendency and personalised therapeutics.

## Introduction

Platelets play a central role in haemostasis but can also acquire a pro-thrombotic phenotype predisposing to ischaemic events, notably stroke and myocardial infarction, which are major causes of mortality and morbidity worldwide^1–3^. Thrombosis risk is based upon multiple factors. Firstly, vascular dysregulation and interventions such as stent and valve implantations. Second, it is increasingly evident that inflammation produces an interplay between platelets and immune cells in a process called thromboinflammation which is a common element in the pathophysiology of cardiovascular disease^4,5^. The COVID-19 pandemic exemplified this with patients progressing to respiratory failure characterised by thrombosis throughout the lung^6–8^. Beyond infection, thromboinflammation is implicated in a breadth of disease scenarios^6^, including cancer, diabetes, vascular dementia and arthritis amongst others. This indicates the widespread importance of identifying patients with hyper-reactive platelets to inform patient care. However, current platelet function tests to measure individualised risk are inadequate and are not routinely used in practice. This highlights the need for new, more effective platelet function tests, for assessing patient-level thrombotic tendency.

Platelet activation is typified by a variety of single platelet and system-level characteristics that have received attention as biomarkers of platelet function. Numerous instruments^10^ have been used to detect these biomarkers, ranging from flow cytometry for the detection of surface markers^11,12^, to light transmission aggregometry^13^ or impedance-based systems^14^, and systems measuring the biomechanical properties of clot formation ^15^ such as thromboelastography^16^. While each have been shown to have value for measuring functional deficits allowing the diagnosis of inherited platelet disorders or measuring the efficacy of anti-platelet therapy, they cannot offer a truly individualised assessment of platelet-mediated thrombotic risk. Despite the high clinical value, platelet function tests have not fundamentally changed in decades, such that the need remains for new assays and biomarkers to identify patients with thrombosis risk.

The concept of platelet functional heterogeneity is long established^17–20^, beginning with considerations of age and metabolic decline^21^, and then size^22–24^ and structural properties^25^. From these seminal findings emerged the appreciation of immature or reticulated platelets, newly formed platelets that are larger and with higher RNA content^26^, higher reactivity^16,27^ and linked to stroke^28^, acute coronary syndrome^26,30^ and cardiovascular disease in general^31^. While platelet lifespan is delimited by apoptosis^32,33^, ageing and functional decline^34,35^ represents a continuum^16^, making cutoffs necessarily arbitrary^36^. In addition, platelet adaptation in disease and to their microenvironment^17,18^, may indicate the need for other measures, beyond size and RNA content, to define pathological platelets.

Beyond reticulated platelets, the topic of platelet functional heterogeneity has received continued interest, with the definition of several platelet sub-populations with functional specialisations^17,16,37–41^. To better understand platelet specialisations, their role in disease, diagnostic possibilities and approaches to treatment new analytical methods are needed^5,18^. This necessarily requires techniques allowing the high throughput analysis of single platelets. Microfluidics using droplets^42^ or microfabricated compartments^43^ has become a powerful high throughput single-cell analysis approach, with examples including the analysis of genomes^44,45^, epigenomes^46^, transcriptomes^47,48^, proteomes^46,50^ and metabolomes^51^. To service the immunology field, microfluidic platforms have been developed for the measurement of single-cell secretions^52–57^. In the context of platelet function testing, multiple platelets have been packaged within large droplets to measure the clotting time^58^ or propensity to aggregate^56,60^, but the ensembled responses neglect single-platelet heterogeneity.

To measure platelet functional variation, we previously developed a single platelet droplet microfluidics and flow cytometry workflow, revealing a sensitivity continuum and an intrinsic heterotypic polarisation program^61,62^. Taking new directions, in this study we combined droplet microfluidics with imaging for the fluorescent detection of ADP secretions using a commercially available reporter. Of note, ADP secretion is a hallmark of platelet activation and underpins the amplification mechanism for platelet cooperation. Illustrating the importance of ADP secretion, numerous anti-platelet medications target the P2Y_12_ ADP receptor. Confining ADP secretions with droplets enhances assay sensitivity and with imaging as a readout allows the high throughput profiling of platelet heterogeneity. In contrast, current luminescent methods for the detection of ATP, another dense granule secretion, involve the ensembled analysis of many thousands of platelets, overlooking platelet heterogeneity. With the convenience of ADP analysis and the benefits of droplet microfluidics the method was used to distinguish resting, primed, activated and hyper-active platelet states, and further used to assess responses to aspirin and P2Y_12_ inhibitors. This approach has potential to define individual tendency to platelet-mediated thrombosis and thereby provide a logical platform for therapeutic targeting.

## Results and Discussion

### The Droplet ADP Secretion Assay

ADP secretion is a characteristic of platelet activation, propagating activation to provide a system-level amplifier in response to stimulation^63^. These features make ADP secretion an appealing biomarker with potential to indicate thrombosis risk. Droplet containment acts to confine ADP secretions to increase assay sensitivity. Droplets also afford the possibility for single platelet encapsulation to allow platelet functional heterogeneity to be surveyed. Microfluidic devices (Figure 1A) were used to encapsulate platelets in 3.3 pL monodisperse droplets along with a fluorescent ADP reporter. Stimulation with 10 ng/mL convulxin resulted in red fluorescent droplets, representing those containing platelets secreting ADP (Figure 1B). The method is termed the Droplet ADP Secretion Assay (DASA). To evaluate the sensitivity and dynamic range of DASA a droplet ADP standard curve was prepared. DASA was quantitative with a limit of detection of 500 nM (1.5 μM limit of quantitation) and a dynamic range of 0.5–30 μM (Figure 1C).

**Figure 1.**
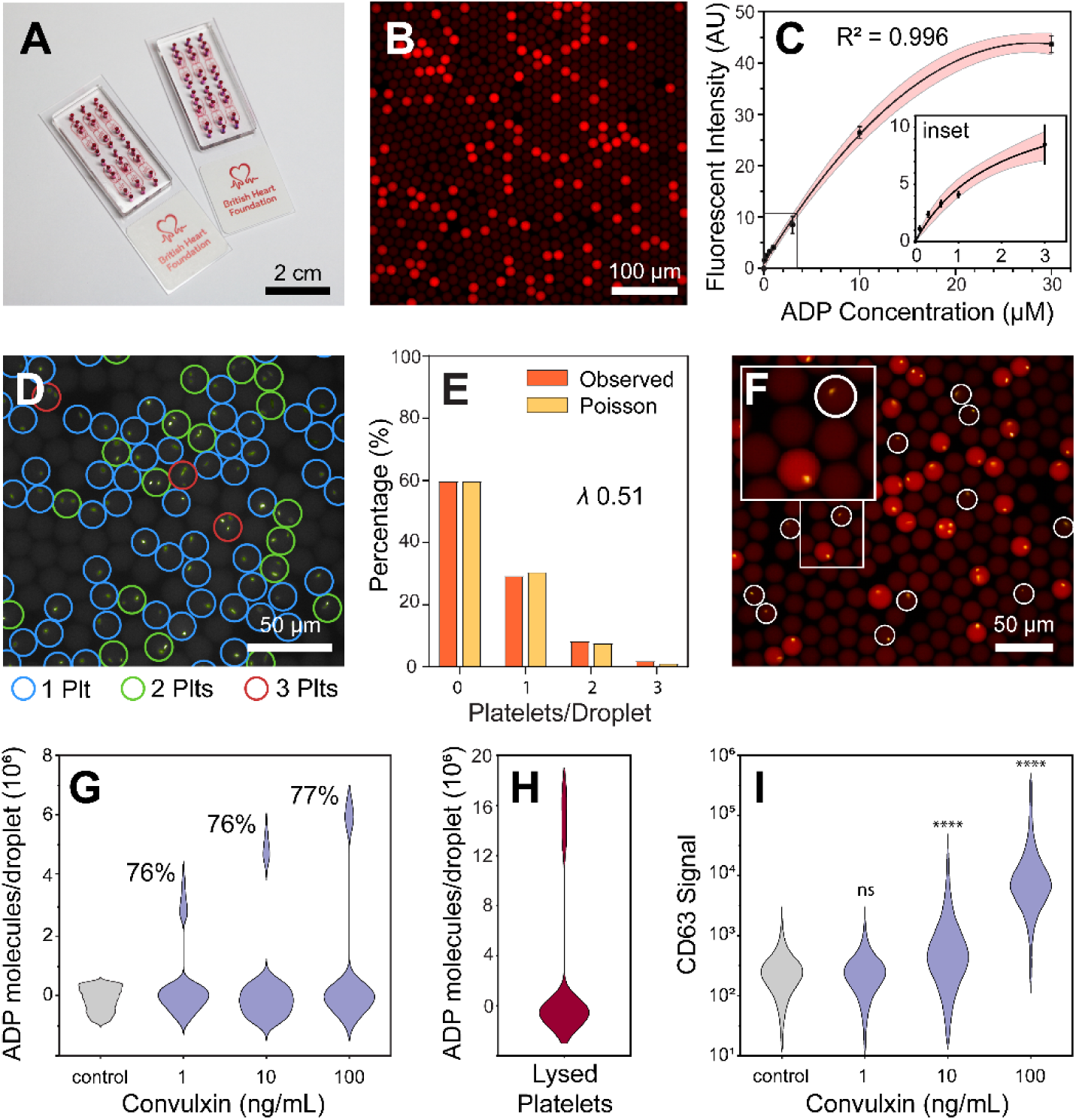
*The Droplet ADP Secretion Assay (DASA).* Array of 6 droplet microfluidic devices on each microscope slide to facilitate multi-sample processing (A). A red fluorescent reporter allows the detection of ADP secreted by single platelets in 3.3 pL droplets (B). ADP can be quantified with a limit of detection of 500 nM (C). Droplet occupancy follows a Poisson distribution (D,E). Platelets can be stained with Calcein AM, unambiguously showing heterogeneous ADP secretion (F). Dose dependent ADP secretion in response to convulxin, with ∼3/4 of platelets from this donor responding (G). Platelet lysis releases ADP from dense granules and other compartments, revealing a large ADP reservoir (>10^7^ molecules/platelet, H). Flow cytometry convulxin dose response using CD63 as the end point (I).

The frequency of platelet encapsulation was investigated using Calcien AM as a live cell dye. Encapsulation followed a Poisson distribution in accordance with the platelet-rich plasma (PRP) concentration. In typical experiments a 2 × 10^8^/mL platelet input into 3.3 pL droplets with 3-fold dilution with reagents produces a mean platelet per droplet occupancy (*λ*) of 0.23, with 80% droplets being empty, 18% of droplets containing single platelets and 2% containing two or more platelets. The Poisson distribution was validated using different input concentrations of PRP (SI Figure 1). The encapsulation data indicates all platelets are delivered to droplets without losses by sedimentation^64^ or adhesion to tubing or microchannel surfaces (Figure 1D,E, SI Figure 1). In Figure 1D, a high lambda value (*λ* 0.51) was used to aid visual illustration. Combining signals from Calcein and the ADP reporter it was evident that ∼1/4 of viable platelets did not secrete ADP on challenge with convulxin, revealing a digital heterogeneous response (Figure 1F). The non-secreting platelets may be aged, and in an exhausted state^65^.

DASA measurements of ADP secretion were reproducible, with equivalent responses to convulxin (collagen substitute and potent GPVI-specific receptor agonist^66^) in a replicate experiment with PRP from the same donor (SI Figure 2). We next investigated the tunable nature of ADP secretion. Responses to 1–100 ng/mL convulxin treatments were dose-dependent, ranging from ∼3 million ADP molecules per platelet to ∼6 million (values obtained by reference to a droplet ADP standard curve prepared in parallel). In contrast the fraction of responding platelets was unaffected (Figure 1G), indicating the same platelets, irrespective of convulxin dose, secreted ADP. The sizeable lower bulb of the violin plots reflects the large number of empty droplets (as well as droplets containing non-secreting platelets). Droplet redundancy is required for reducing multiple platelet loading necessary for single platelet anaysis, albeit with a few droplets (∼2%) containing multiple platelets to favour assay throughput. The low signal from empty droplets and those containing non-secreting platelets results from the millisecond timescales^67^ between platelet encounter with agonists and droplet encapulation, a time insufficient for ADP secretion, dispersion and contamination within other droplets.

**Figure 2.**
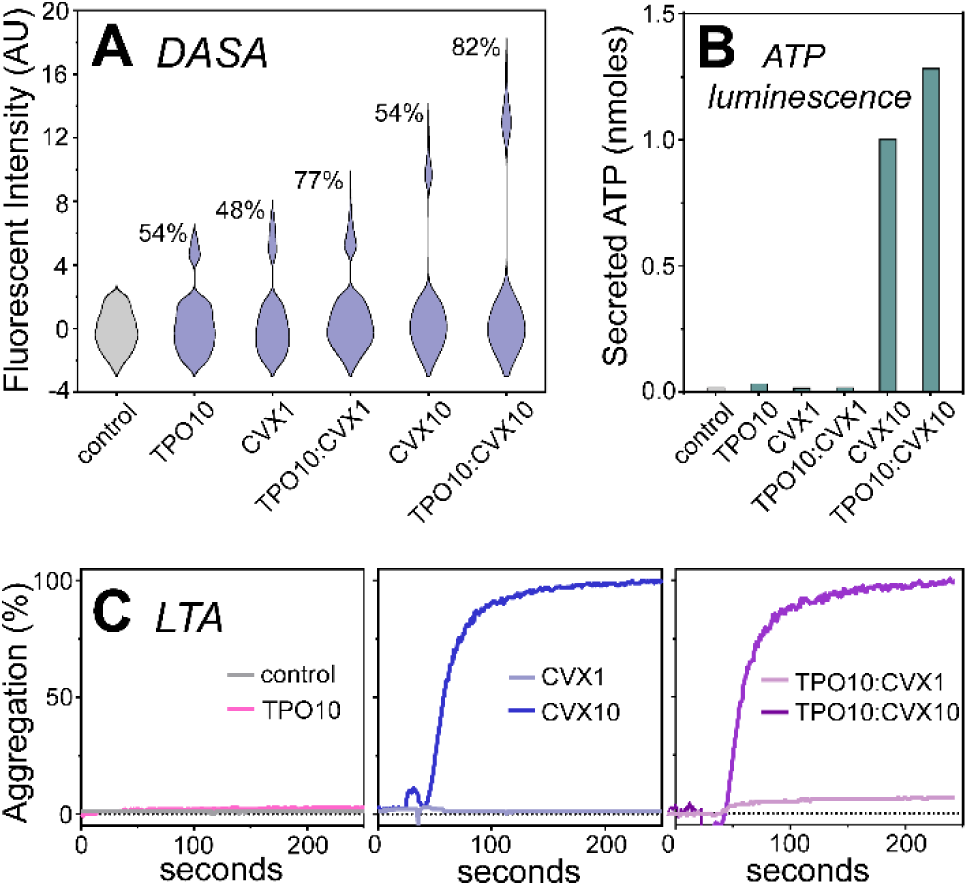
*Benchmarking DASA against bulk platelet function tests.* Platelets were treated with convulxin (CVX, 1 or 10 ng/mL) with or without thrombopoietin (TPO, 10 μg/mL). DASA produces signals for all treatments (A), whereas only treatment with 10 ng/mL convulxin, with or without TPO priming, produced distinct responses from an ATP luminescence assay (Chrono-Log 700, B) or light transmission aggregometry (Helena Biosciences, C).

To appreciate the available ADP pool within platelets thermal platelet lysis was used to release ADP stores from dense granules, and cytosolic and mitochondrial compartments. Approximately 16 million ADP molecules per platelet are available indicating capacity for tunable and successive secretions (Figure 1H). ADP quantification requires a standard curve, with subsequuent experiments only reporting droplet fluorescent intensity values. Cytometry measurements of CD63 presentation, an alternative marker of dense granule secretion, also revealed a dose-dependent response albeit only with higher convulxin concentrations of 10 ng/mL and 100 ng/mL (Figure 1I). P-selectin exposure, a marker of alpha granule secretion, is known to be a more sensitive and robust indicator of platelet activation than CD63^68^. This was confirmed in a convulxin dose response experiment. Despite gains over CD63, P-selectin was still 10-fold less sensitive to platelet responses compared to DASA (SI Figure 3).

**Figure 3.**
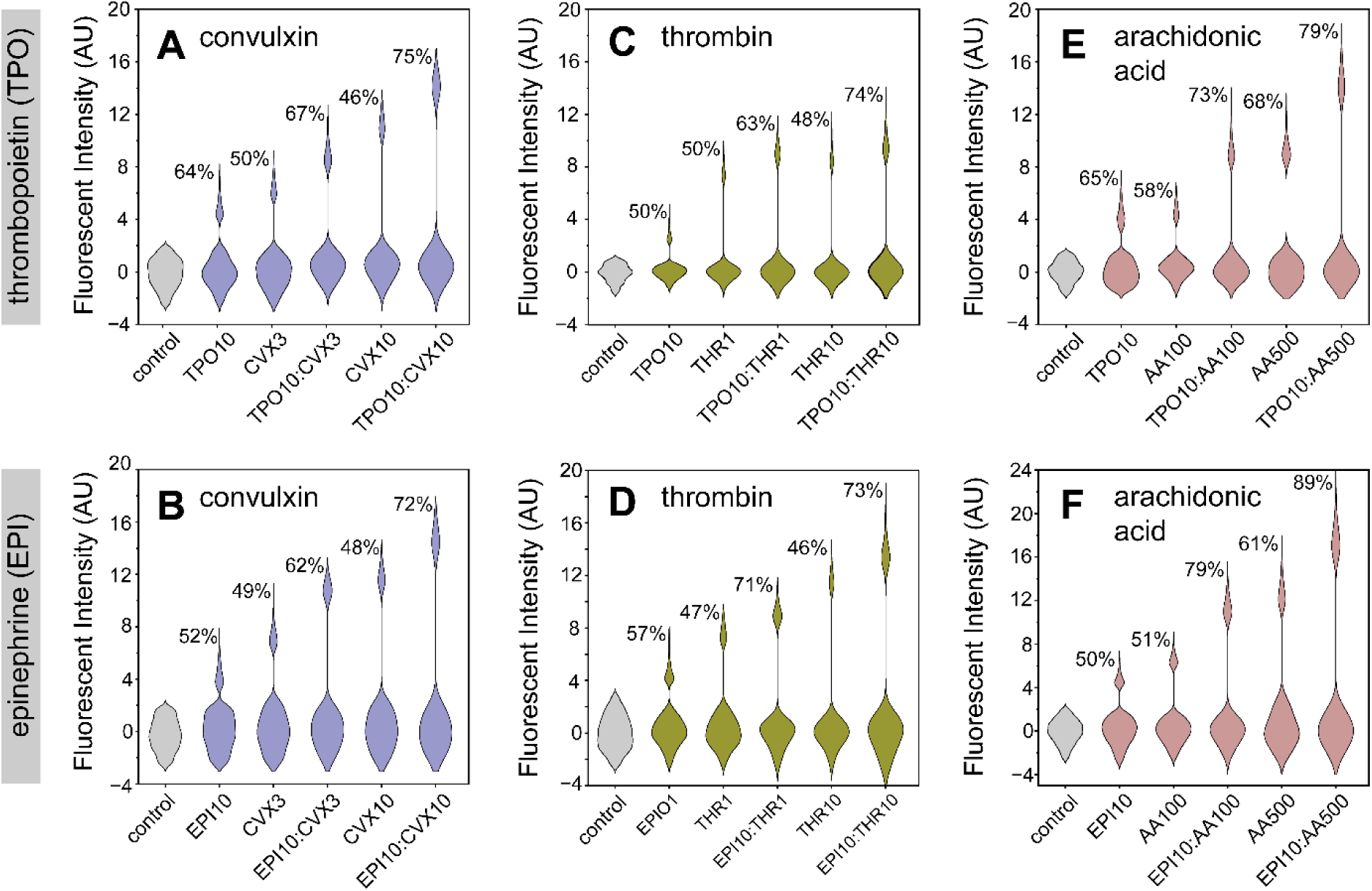
*ADP secretion generality and synergy between agonists and primers.* Platelets were stimulated with convulxin (CVX, lilac, 3 or 10 ng/mL, A,B), thrombin (THR, green, 1 or 10 IU, C,D) or arachidonic acid (AA, pink 100 or 500 μM, E,F) agonists alone or following priming with thrombopoietin (TPO, 10 μg/mL) or epinephrine (EPI, 10 μM). Control intensity distributions are in grey. Dose-dependent ADP secretion responses were observed along with primer-agonist synergy increasing ADP secretion levels and the fraction of secreting platelets.

While CD63 presentation and ADP secretion both report dense granule release and both DASA and flow cytometry (CD63 or P-selectin) employ fluorescent detection principles, the difference in sensitivity deserves consideration. DASA has tight data distributions (coefficient of variation, CV<10%) supported by the high degree of droplet monodispersity (CV<4%). In comparison, flow cytometry data is broadly distributed (CV∼90%) increasing the baseline defining the limit of detection. In addition, DASA sensitivity is aided by secretions being confined within droplets, effectively concentrating ADP to increase the fluorescent signal.

### Priming and Benchmarking

Platelet priming is known to potentiate platelets, producing elevated sensitivity and responses to agonists, and is implicated in thromboinflammation and resistance to anti-platelet medication^17,69,70^. Experimentally, priming is commonly observed by dual treatment with a primer and an agonist, producing a higher response than treatment with agonist alone. We applied DASA to investigate platelet responses to priming by thrombopoietin (TPO, 1–100 μg/mL)^71^ and epinephrine (EPI, 1–100 μM)^72^. Of significance, primed platelets secreted ADP in the absence of agonists. The level of ADP secretion was dose dependent without markedly affecting the fraction of responding platelets (SI Figure 4A,D). In contrast, flow cytometry measurements involving CD63 and P-selectin markers did not, as expected, produce signals above the vehicle control, excepting treatment with 100 μM EPI (SI Figure 4B,C,E,F), consistent with EPI being a weak agonist^73^. The absence of CD63 signals in response to priming has been described by others^74^. While DASA may have higher sensitivity than flow cytometry, it does not exclude the possibility that priming could cause ADP to be released from stores that do not coincide with CD63 presentation.

**Figure 4.**
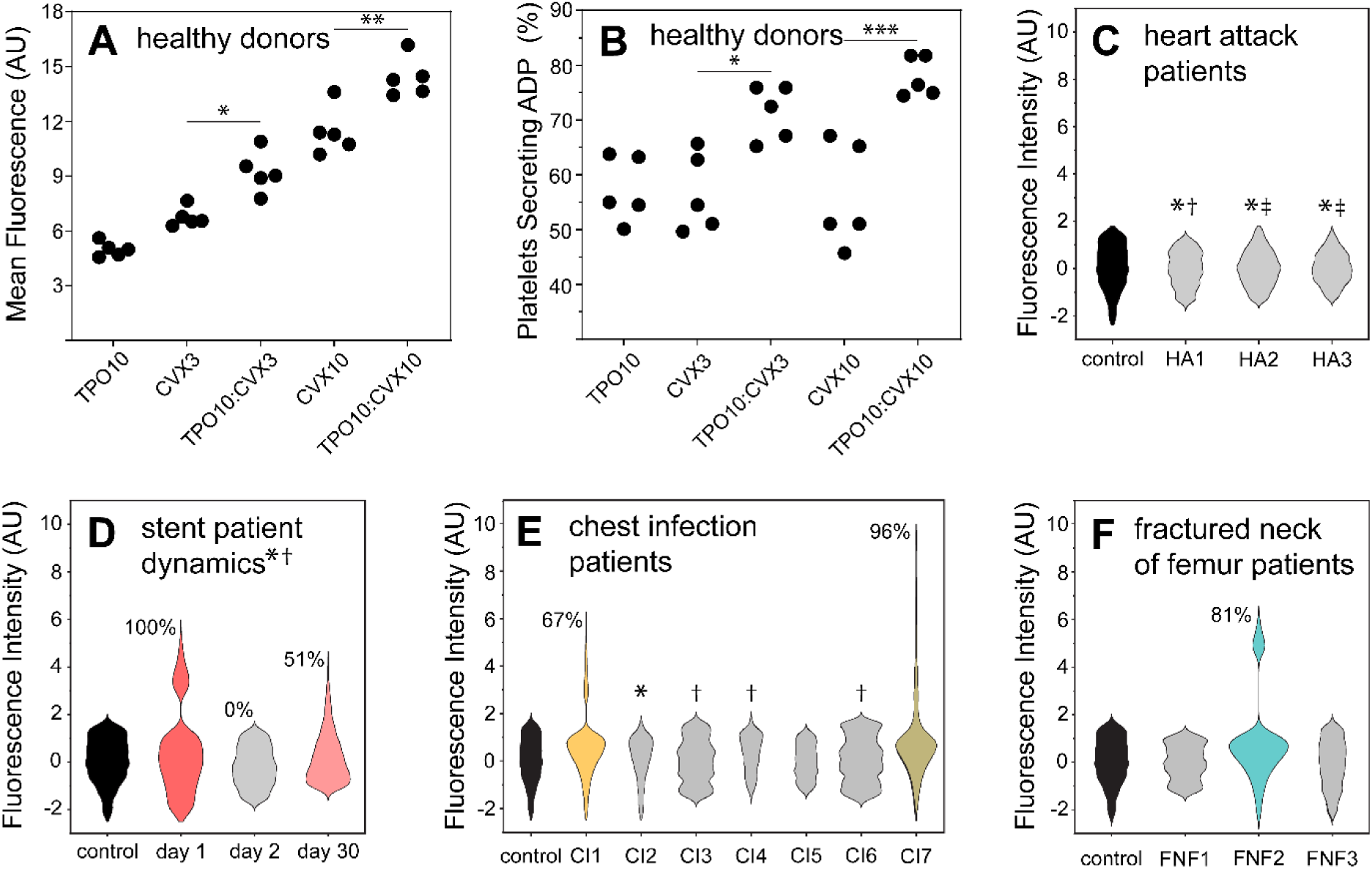
*Putative health and risk DASA signatures.* ADP secretion levels (A) and fraction of secreting platelets (B) for five different healthy donors (age, gender and contraceptive). Platelets from healthy donors were treated with convulxin (CVX, 3 and 10 ng/mL), thrombopoietin (TPO, 10 μg/mL) and dual treatment. Significance determined using a one-way ANOVA with Tukey’s test. Platelets from patients were not challenged with agonist, with ADP secretions used to infer *in vivo* status. DASA results of platelets from heart attack patients, receiving aspirin* and clopidogrel^†^ or prasugrel^‡^, (C). 30-day monitoring of a patient admitted for acute myocardial infarction due to stent thrombosis and receiving aspirin* and clopidogrel^†^ (D), chest infection patients with indicated patients receiving aspirin* or clopidogrel^†^ (E) and fractured neck of femur patients (F).

Increased responses to agonists by primed platelets are known from the analysis of different biological features such as expression of surface markers measured by flow cytometry and aggregation measured by light transmission aggregometry. With DASA, TPO priming and stimulation with convulxin also produced a synergistic effect, increasing the level of ADP secreted and the fraction of responding platelets. This demonstrates that additional platelets can respond, converted from an otherwise inactive state by priming (Figure 2A). The finding that non-responsive platelets can progress to secrete ADP when activation pathways are potentiated by priming, indicates this platelet subpopulation has pathway deficits, not deficits with dense granules or their contents.

The higher sensitivity of DASA than flow cytometry, both single platelet analysis methods, prompted benchmarking against conventional bulk assays. The ATP luminescence assay also reports dense granule secretion, but unlike DASA did not produce measurable signals in response to 10 μg/mL TPO or 1 ng/mL convulxin. The assay did produce signals in response to 10 ng/mL convulxin along with a gain in signal when also treated with TPO (Figure 2B). Here we attribute the higher sensitivity of DASA, able to detect responses to priming and 1 ng/mL convulxin, to ADP confinement within droplets, but also the higher background signal associated with bulk assays.

Moving to light transmission aggregometry (LTA), widely regarded as the gold standard platelet function test, DASA again had higher sensitivity, with LTA unable to detect responses to TPO or 1 ng/mL convulxin. LTA did, as anticipated, produce a signal, albeit minor, in response to 1 ng/mL convulxin when primed with TPO, and robust aggregation when stimulated with 10 ng/mL convulxin alone or along with TPO priming (Figure 2C). While the sensitivity of DASA allows responses to primers and low agonist doses, these features do not correlate with LTA measurements of aggregation. While ADP secretion by primed platelets does not cause aggregation *in vitro*, such experiments involve only a small fraction of the >1 trillion platelets in circulation. Such numbers combined with multiple sites of even minor vessel damage or other platelet triggers, introduces potential for localised platelet activation, the sum of which may indicate thrombosis risk. Testing this hypothesis is beyond the reach of experimental methods, instead probabilities can be obtained using mathematical methods^75^. We envisage the ability to measure ADP secretions from primed platelets, a state predicting increased reactivity on future exposure to agonists and causing system amplification, indicates potential to use DASA as a snapshot early warning system to alert clinicians to elevated thrombosis risk.

### Generality, and Healthy and Risk Signatures

To explore the generality of ADP secretion as a marker of platelet reactivity a screen with other agonists and priming agents was undertaken. Convulxin^76,77^, thrombin^78^ and arachidonic acid^76^ agonists and the primers thrombopoietin^80^ and epinephrine^81^ have distinct receptors and different signalling pathways, with some converging more than others^82,83^. Nevertheless, DASA reported consistent trends, detecting primed states, dose responses to the different agonists and synergy with combined primer and agonist treatment (Figure 3). In all cases, synergy produced increases in ADP secretion levels and the responding platelet fraction, lending weight to the argument of priming restoring functionality in a subset of otherwise inactive platelets.

The same samples were also measured by flow cytometry using the P-selectin endpoint (improved sensitivity compared to CD63) did not detect the primed states or report clear dose responses to agonists (SI Figure 5). Synergy between primers and agonists was not observed except in the case of platelets treated with 3 ng/nL convulxin in which priming with epinephrine converted platelets from a resting to an activated state (SI Figure 5B). These data provide a first indication of the broader potential of DASA to measure platelet responses to different agonists and in different priming contexts. However, the DASA readout necessarily precludes the use of ADP as an agonist.

**Figure 5.**
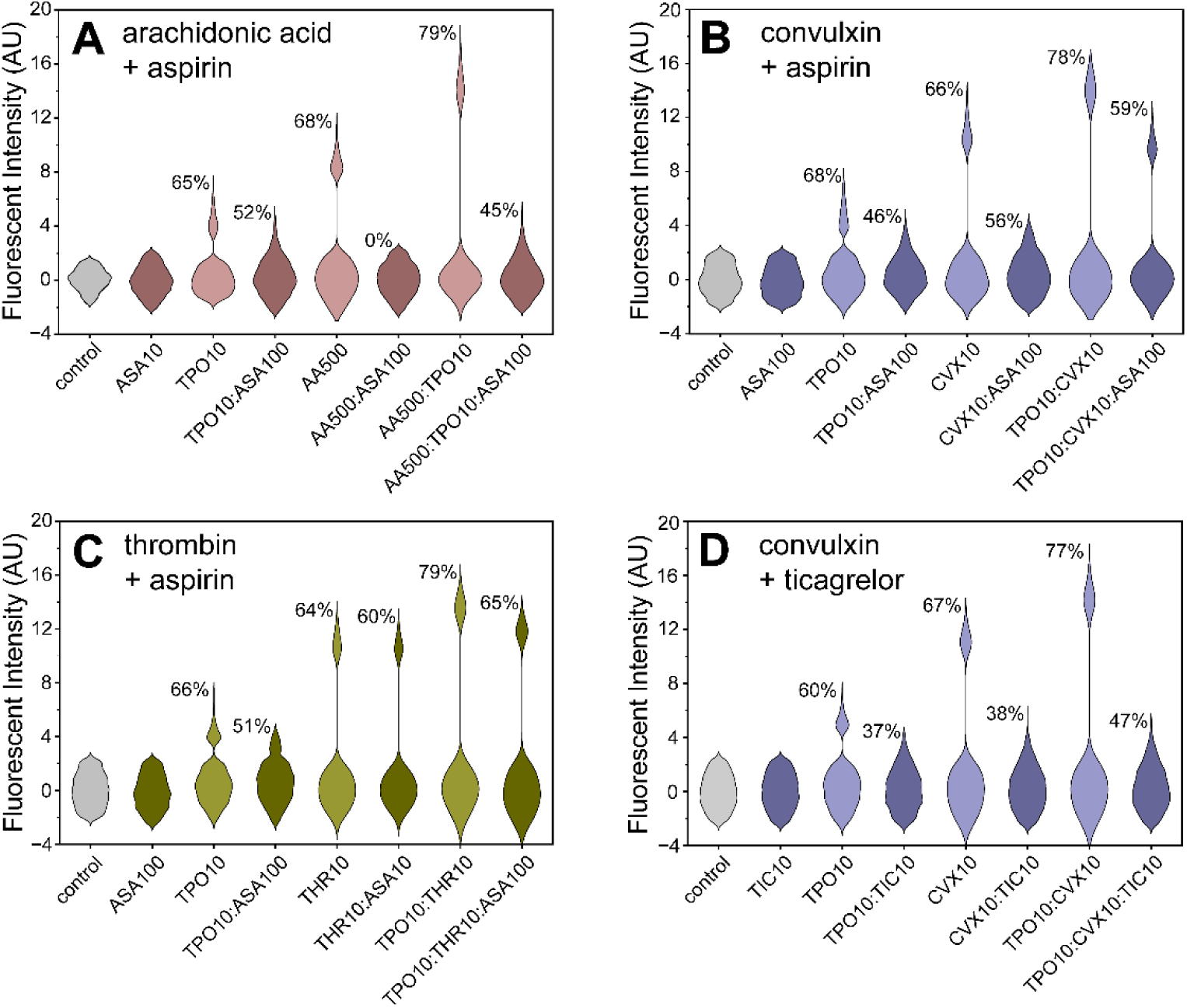
*DASA measures of response to anti-platelet therapy.* Treatment with 100 μM aspirin (ASA) reduced ADP secretion levels and fraction of ADP secreting platelets in platelets primed with thrombopoietin (TPO, A–C), and platelets treated with arachidonic acid (AA, pink, A) and convulxin (CVX, blue, B). Aspirin did not alter ADP secretion responses to thrombin (THR, green C). Increased ADP secretion in response to dual treatment with TPO and agonists (AA, CVX and THR) is prevented by aspirin (A–C). Treatment with 10 μM ticagrelor (TIC) reduced ADP secretion levels and secreting platelet fraction responses to TPO priming, CVX stimulation and dual treatment with TPO and CVX (D).

The precision DASA datasets were used to evaluate whether a platelet reactivity signature could be described for healthy donors. DASA results of platelets from 5 healthy donors, male and female between 20 and 50 years of age were obtained (SI Figure 6). ADP secretion levels were tightly grouped (CV 8–13%) within agonist, primer and dual treatment conditions, as were the fraction of secreting platelets (CV 5–17%) (Figure 4A,B). To understand platelet dynamics, we measured the reactivity of platelets sampled from a young (25) healthy female donor over a 4- month period. Again, ADP secretion levels and fraction of responding platelets were tightly grouped, albeit with one exception. On one occasion there was a high responding fraction (77% vs ≤50%) of platelets to TPO, which may coincide with menstruation^84^. Whether DASA metrics truly represent a signature of platelets in health providing a baseline to discern disease requires much larger and broadly inclusive investigations to ascertain clinical value.

**Figure 6.**
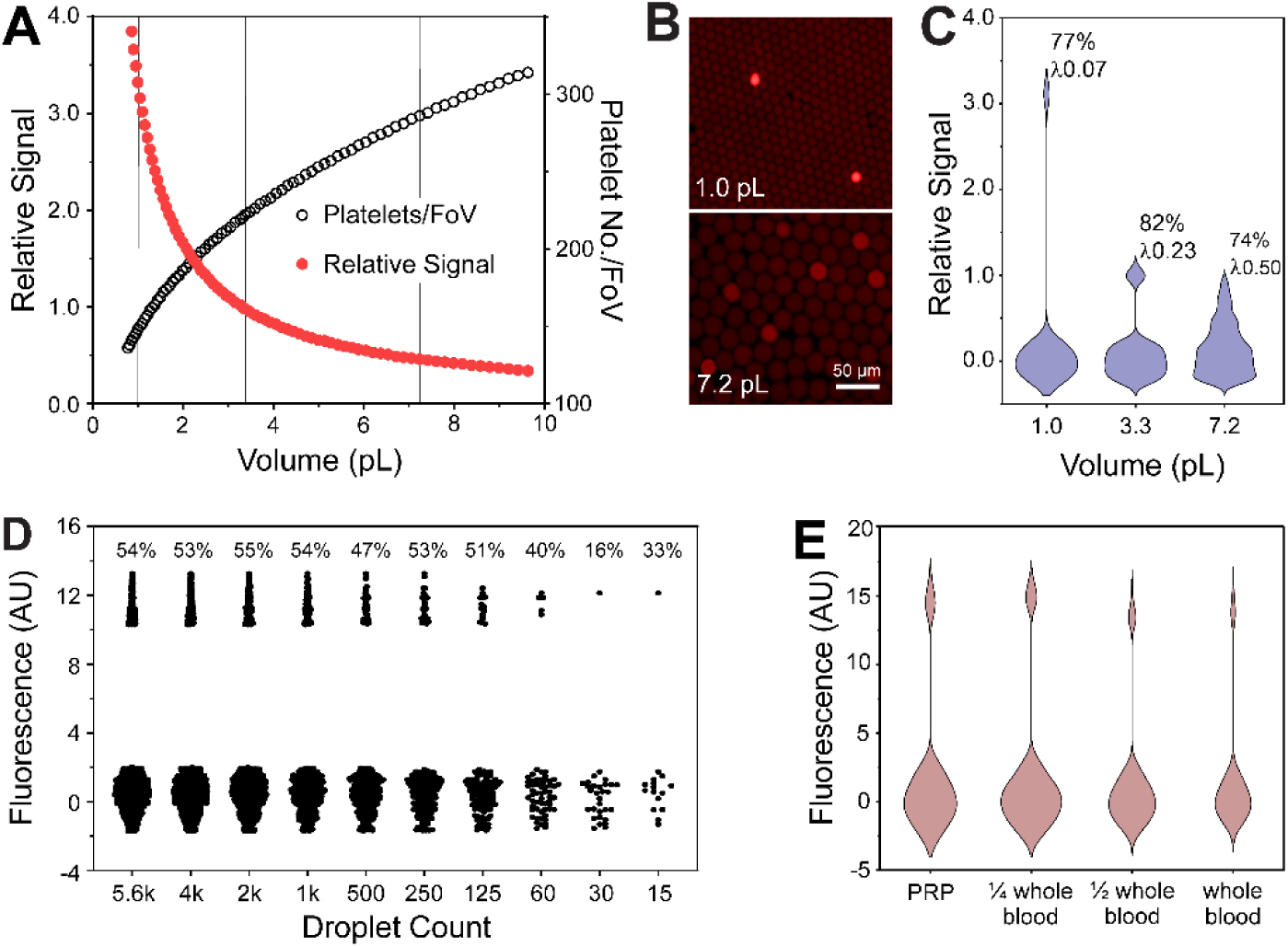
*DASA refinement.* Theoretical optimal droplet volume in terms of sensitivity and platelet number in the field of view (FoV, A). Droplet miniaturisation from 7.2 to 1.0 pL increases sensitivity proportionately and reduced the mean platelet per droplet occupancy (λ) from 0.50 to 0.07 (B,C). Randomised data reduction used to estimate minimal data requirements for accurate measurements (D). Comparison of DASA results from platelet-rich plasma and whole blood dilution samples (E).

As a first step towards clinical observations, platelet samples were obtained, on occasion, from a small clinical study involving different patient cohorts at risk of thrombosis. Platelets were measured without agonist challenge, to provide an indication of their recent *in vivo* state (*i.e.* are platelets primed?). ADP secretion was not detected in the three patients from the acute myocardial infarction following inflammation patients (Figure 4C). These patients were receiving aspirin and either prasugrel or clopidogrel (P2Y_12_, ADP receptor inhibitors), likely indicating these medications are effective. Platelets from a patient with acute myocardial infarction due to stent thrombosis were monitored (Figure 4D). On admission, all platelets were secreting ADP at levels typically seen with primed platelets from healthy donors. This indicates that although aspirin and clopidogrel loading doses had been administered they had yet to take full effect. By day two, platelets had ceased secreting ADP. Analysis on day thirty showed that despite aspirin and clopidogrel maintenance almost half of the platelets were secreting ADP albeit at low levels, potentially indicating transition to a risk phase.

In the context of chest infection, two of seven severe chest infection patients had platelets secreting ADP, one of which had 96% of platelets secreting ADP (Figure 4E), suggesting these patients had platelets in a primed state. Of note, four of the five patients without ADP-secreting platelets were receiving either aspirin or clopidogrel, which may indicate their protective benefits. In the fractured neck of femur cohort, one of three patients had platelets secreting ADP, an interesting observation given the association between fractured neck of femur and subsequent acute MI^85^ (Figure 4F). The data suggests that ADP secretion in the absence of agonist stimulation, may represent a useful biomarker for the identification of patients at risk. Beyond this, the level of ADP secretion, the concentration of platelets and secreting fraction may provide risk metrics. Patients with high values may warrant treatment with P2Y_12_ inhibitors to ameliorate the effects of ADP secretion.

For comparison, all patient samples were also analysed using a convulxin dose-response experiment using the P-selectin biomarker measured by flow cytometry (SI Figure 8). Results were ambiguous, with patient platelet reactivity lower than controls from healthy donors, perhaps indicating exhausted platelets^65^. In summary, the cytometry data using P-selectin as an activation marker is less suited for identifying patients with elevated platelet reactivity. Alternatively, commercial cytometers (*e.g.* Mindray, Sysmex and Cell-Dyn Sapphire) can be used to detect reticulated platelets based on size and elevated RNA content. Reticulated platelets have high reactivity and are likely to be ADP secretors. Given the majority fraction of platelets in a sample are ADP secretors, there may be merit in extending the reticulated platelet cut-off^36^ with the onus on excluding aged platelets to provide a more accurate risk score.

We next investigated whether DASA could measure responses to aspirin the frontline anti-platelet medication^86^. Aspirin blocks cyclooxygenase-1, preventing arachidonic acid being converted to thromboxane A2, inhibiting response to arachidonic acid, and inhibits ADP secretion responses to thrombopoietin priming, by removing the amplification effects of thromboxane A2 (Figure 5A). Aspirin also inhibited ADP secretion responses to convulxin^87^, indicating the importance of thromboxane A2 for full response to agonism (Figure 5B). However, aspirin only elicited a minor reduction in ADP secretion and fraction of responding platelets when platelets were also primed with thrombopoietin. This indicates the ability of platelet priming to override the effects of aspirin. With thrombin stimulation, aspirin was unable to inhibit responses to the potent agonist thrombin^88^, while aspirin did reduce priming effects (Figure 5C). Moving to ticagrelor, a P2Y_12_ (ADP) receptor inhibitor and not requiring metabolic conversion, convulxin activation and thrombopoietin priming effects were almost completely abolished (Figure 5D), illustrating the importance of ADP autocrine feedback involvement in platelet activation. The ability to measure response to anti-platelet treatment may prove valuable for personalised medicine, for example in order to identify those patients unresponsive to conventional medications and needing a bespoke management plan^89–91^.

Given the merits of DASA for detecting different platelet response states and provide activation signatures from at-risk patient samples coupled with potential to indicate medication suitability we investigated routes to optimisation. First, theory was used to optimise droplet volume. Droplet miniaturisation concentrates ADP to increase sensitivity while reducing mean platelet droplet occupancy (lambda), albeit with more smaller droplets displayed in the 600-μm- diameter field of view. Sensitivity scales linearly with volume, with 3-fold sensitivity gains when moving from 3.3 pL to 1.0 pL volumes, with >100 platelets within the field of view (Figure 6A). Experiments confirmed ADP sensitivity dependence on droplet volume (Figure 6B,C). We next investigated data minimalism. In this example, measurement of as few as 14 platelets was sufficient to provide accurate measures of ADP secretion and platelet fractional response (Figure 6D).

Overall, the data implies droplet miniaturisation affords the possibility of using less sensitive, less costly cameras. For point of care application, DASA should also deliver results in short timescales and involve minimal sample preparation. The fluorescent reporter for ADP has the signal fully developed in 10 minutes (SI Figure 9). Added to this, DASA measurements are unaffected when processing whole blood samples (Figure 6E), bypassing the need for the time-consuming and cumbersome task of preparing platelet-rich plasma. Data minimalism, less costly cameras, rapid result delivery and removal of platelet sorting are attractive for reducing instrument and workflow complexity. DASA is a candidate test to broaden our ability to define individualised platelet-mediated thrombotic risk, having potential in a range of academic and clinical applications, such as personalised approaches to anti-platelet therapy of near patient thrombotic risk assessment. For instance, DASA has potential to analyse a finger prick blood sample with image collection, analysis and reporting to healthcare professionals using a smart phone.

## Conclusions

DASA provides a high throughput and sensitive means to profile single-platelet activity. Digital and tuneable responses were reported across a range of primers and agonists, with synergy leading to increased ADP secretion and the fraction of ADP-secreting platelets. Primed platelets, potentiated for elevated reactivity were shown to secrete ADP. Similar signatures were observed in a variety of patients at risk of thrombosis which may indicate these patients warrant treatment with anti-platelet medications. DASA also reported responses to aspirin and P2Y_12_ inhibitors.

Overall, DASA holds promise for individual patient risk stratification and a personalised medicine approach to therapy selection.

## Materials and Methods

### Participants, Sampling and Treatments

Blood samples were obtained from healthy volunteers after obtaining informed consent under institute (ERGOII: 62658.A1) and South Central-Hampshire B (REC: 22/LO/0801) ethical approvals. Participants were free from anti-platelet medication, such as aspirin for 2 weeks and >24 h free from other non-steroidal anti-inflammatory medication. Venepuncture with a 21G needle was used to collect blood in 8.5 mL vacuum tubes containing ACD (sodium citrate: 22.0 g/L; dextrose: 24.5 g/L; citric acid: 8.0 g/L; antifungal reagent (potassium sorbate: 0.15 g/L) (first 4 mL discarded). Tubes were gently inverted, centrifuged at 220*g* for 12 min without brake to prepare platelet-rich plasma (PRP) that was rested for 30 min prior to experiments. Platelet counts were determined by labelling with a CD61 antibody and detection by flow cytometry using an Attune NxT instrument (ThermoFisher Scientific). Platelets were subsequently diluted to a concentration of 2 × 10^8^/mL in HEPES buffer (136 mM NaCl, 2.7 mM KCl, 10 mM HEPES, 2 mM MgCl_2_, 0.1% (w/v) glucose and 1% (w/v) BSA (pH 7.45)).

Experiments involved untreated platelets (vehicle control) or platelets treated with priming agents and/or agonists during droplet generation. Platelets were primed with thrombopoietin (1–10 μg/mL, recombinant human, PeproTech Inc.) or epinephrine (1–10 μM, Bio/Data Corp.) and stimulated with the agonists convulxin (1–100 ng/mL, Enzo Life Sciences), arachidonic acid (100–500 μM, Bio/Data Corp.) and thrombin (1–10 NIH, Hyphen BioMed). In activity suppression experiments, PRP was incubated with 10 μM ticagrelor (Abcam) or 10 μM acetylsalicylic acid (aspirin, Thermo Scientific Chemicals) for 10 min prior to droplet encapsulation. Thrombin stimulation experiments required the preparation of washed platelets using a modified protocol^62^: Prostacyclin (PGI_2_, 75 ng/mL, Abcam) was added to whole blood and PRP prepared as describe above. Next, PGI_2_ (300 ng/ml) was added to PRP with a further centrifugation at 600*g* for 10 min. The platelet pellet was then washed three times with Tyrode’s buffer and re-suspended in HEPES buffer at 2 × 10^8^/mL, and left to rest for 1 hour to reverse PGI_2_ inhibition.

#### Clinical Cohorts

Ethical approvals (REC: 23/SC/0055) were granted for obtaining and testing blood samples from patients. PRP was prepared and used to characterise platelet samples using DASA without agonist and flow cytometry using P-selectin as an endpoint in a convulxin dose response measurement. This study accessed samples from acute myocardial infarction following inflammation patients (heart attack cohort), an acute myocardial infarction due to stent thrombosis patient, chest infection patients and fractured neck of femur patients. Blood samples from the acute myocardial infarction following inflammation were collected and tested 24 hours from admission, with patients having received loading doses and started maintenance with aspirin and the P2Y_12_ inhibitors clopidogrel or prasugrel. Samples were collected from the acute myocardial infarction due to stent thrombosis patient on admission (day 1, following loading with aspirin and clopidogrel), on day 2 and follow up on day 30 the patient was receiving maintenance doses of aspirin and clopidogrel. Some of the chest infection patients were receiving aspirin or clopidogrel. Fractured neck of femur patients were not receiving anti-platelet medication.

### Microfluidics

Droplet microfluidic devices were fabricated in poly(dimethylsiloxane) (PDMS, Sylgard 184) by classical soft lithography using SU-8 on silicon wafers. SU-8 was spin coated to a height of 14 μm, with a channel width of 30 μm and a droplet junction width of 15 μm. PDMS was cured on the SU-8 wafer at 60 °C for 2 hours. Ports for tubing were introduced using 1-mm-diameter biopsy punches (Miltex, Williams Medical Supplies Ltd). Devices were bonded to glass microscope slides using a 30 s oxygen plasma treatment (Femto, Diener Electronic) followed by channel surface passivation using 1% (v/v) trichloro(1H,1H,2H,2H-perfluorooctyl) silane (Merck) in HFE- 7500 (3M Novec). Syringe pumps (Fusion 100, Chemyx) were used to deliver samples and reagents from plastic syringes (Terumo) interfaced with 25 G needles for connecting to the microfluidic ports via medical grade, sterile polythene tubing (0.38 mm ID, 1.09 mm OD, Smiths Medical). To generate 3.3 pL droplets at 15 kHz, a fluoro-oil (QX200™, Bio-Rad) flow rate of 12 μL/min was used coupled with platelet, treatment and ADP reporter flow rates of 1 μL/min were used. Droplet generation was monitored using a high-speed camera (Phantom Miro M310, Vision Research) mounted on an inverted microscope (Olympus, CKX53). Droplets were collected in microcentrifuge tubes for 10 min (∼10 million droplets) and subsequently an aliquot was pipetted into droplet imaging chambers, fabricated by soft lithography as above to 14 μm in height to ensure single layer droplet packing.

### Fluorescent Analysis

Droplets were imaged using a 16-bit Hamamatsu Fusion camera mounted on an inverted fluorescent microscope (CKX41, Olympus) with a 20x/0.45NA objective providing a field of view with a diameter of 600 μm. Fluorescent excitation at 531±20 nm and emission collection at 593±20 nm with exposure for 120 milliseconds was used to image fluorescence from the ADP reporter (Ex/Em 535/587 nm, ab83359, Abcam). Calcein AM (Ex/Em 495/515 nm, Invitrogen), a live cell dye, was used to identify platelets. For each experimental condition, typically >5,000 droplets were measured. A custom Python script (SI Figure 10) was used for automated droplet detection, droplet fluorescent intensity and platelet count measurements. Signal development from 10 μM ADP with the ab83359 reporter was recorded using a fluorescent plate reader (BMG Labtech Clariostar Plus). Total ADP packaged within platelets was obtained by lysing platelets at 90 °C for 5 min followed by measurement using the fluorescent plate reader.

### Flow Cytometry

Flow cytometry (Attune NxT, ThermoFisher Scientific) was used as a single platelet method against which to benchmark the droplet ADP secretion assay. Fluorescent antibodies from BD Bioscience were used to detect platelets (CD42b conjugated to APC (HIP1) and biomarkers: P-selectin was detected using mouse anti-human CD62P conjugated to APC (AK-4), and CD63 was detected using mouse anti-human CD63 conjugated to FITC (PAC-1). For each experimental condition >10,000 platelet events were recorded. A one-way ANOVA with Tukey’s test was used to compare data from treated platelets with vehicle controls.

### Light Transmission Aggregometry and ATP Luciferase Assay

Platelet aggregation assays were performed using a Helena Biosciences AggRAM™ aggregometer, with platelets stimulated by the addition of 10 μL convulxin (1 and 10 ng/mL final concentration) to 390 μL PRP (previously incubated at 37 °C for 2 min) with continuous stirring. For priming conditions, PRP was incubated for 30 minutes at 37 °C with 10 μg/mL thrombopoietin before agonist addition^93^. The ATP luminescence assay entailed the use of a Chrono-Log® 700 instrument, with quantification calibration using an ATP standard (2 nmole). The assay involved preparing a mixture of 350 μL PRP, 40 μL Chrono-Lume reagent (2 μM luciferin/luciferase) with incubation at 37 °C for 2 min, followed by addition of 10 μL convulxin (1 and 10 ng/mL final concentration). For priming conditions, platelets were incubated with thrombopoietin as described for the light transmission aggregometry experiment.

## Acknowledgements

The authors thank the British Heart Foundation (NH/F/22/70010, MS and JW) and Haemonetics Corp. (AE and NC) for funding. The authors thank Hiroki Cook for support with fluorescent microscopy and Johan W. M. Heemskerk for manuscript feedback.

## Data availability

The raw cytometry.csv files are made available via FigShare: 10.6084/m9.figshare.33060650

## Ethics declarations

The authors declare no competing interests.

