## Supplementary Figures for "Droplet microfluidics reports digital and tuneable single-platelet ADP secretion describing primed, active and hyper-active states"

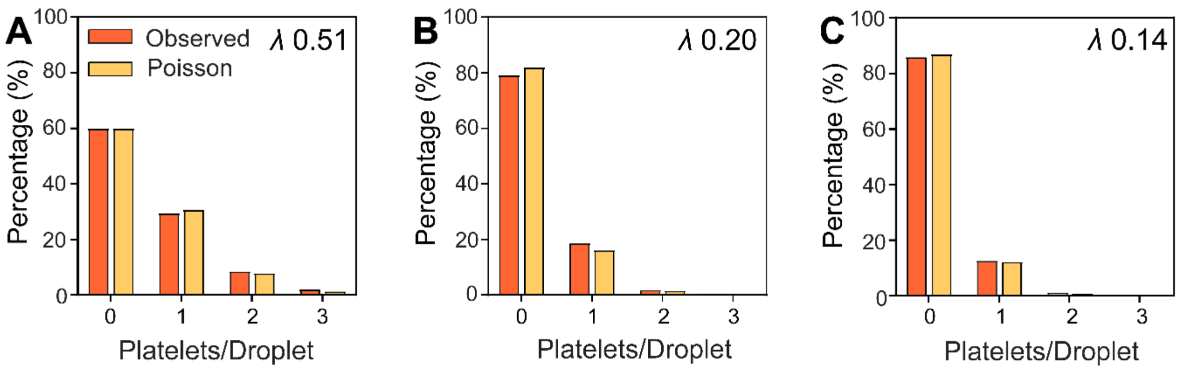


**SI Figure 1.** *Droplet occupancy statistics.* Platelet encapsulation (orange) within droplets closely follows a Poisson distribution (yellow) for tested lambda values of 0.51 (A), 0.20 (B) and 0.14 (C).

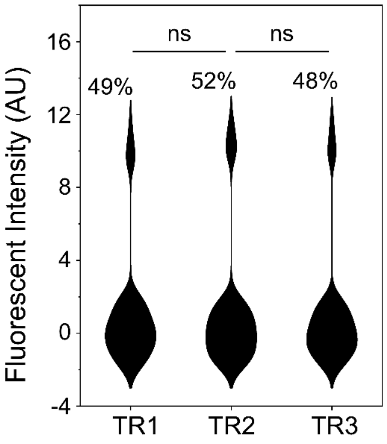


**SI Figure 2.** *Reproducibility.* Three technical replicates (TR) using the same platelet-rich plasma (PRP) sample treated with 10 ng/mL convulxin. Non-significance determined using a two-way ANOVA with Tukey's test.

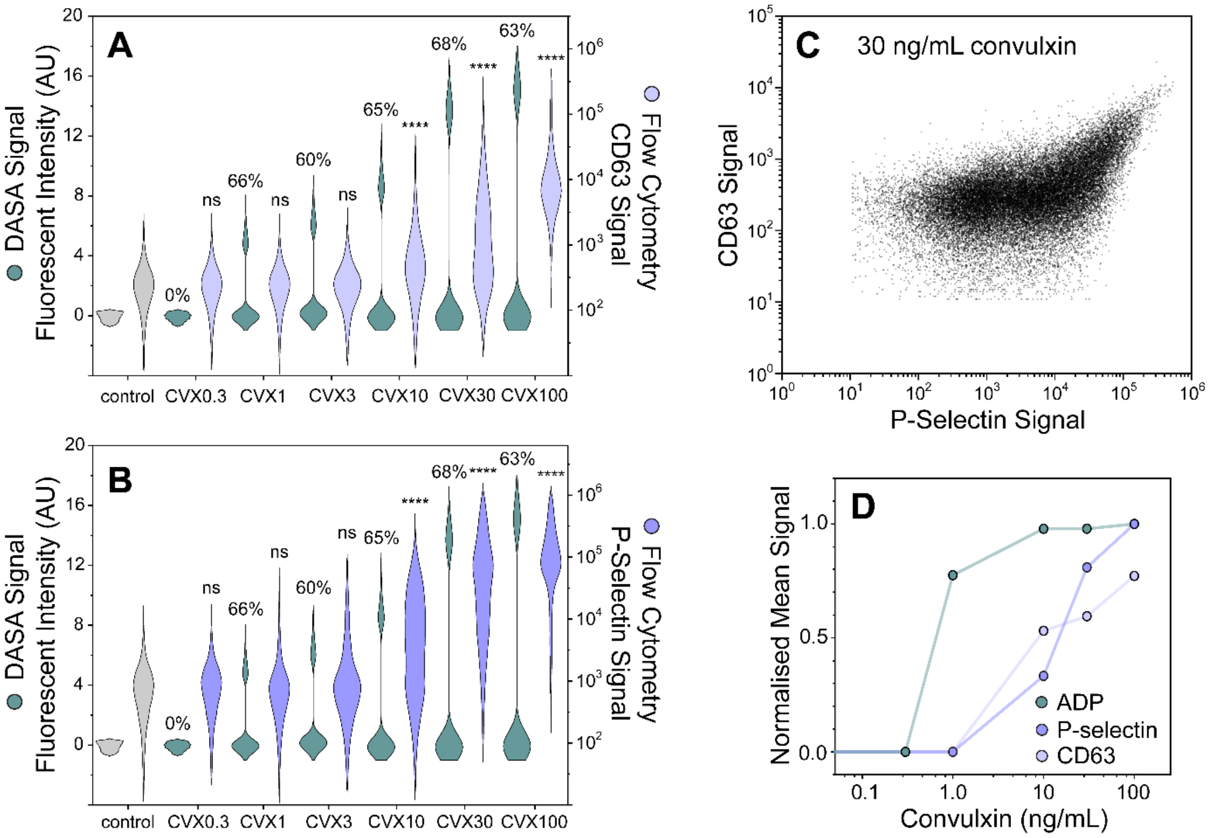


**SI Figure 3.** *DASA sensitivity and dynamic range compared with flow cytometry.* DASA results from a convulxin dose-response experiment compared with flow cytometry data using CD63 (A) and P-selectin (B) endpoints. Significance relative to vehicle controls determined using a one-way ANOVA with Tukey's test. The P-selectin activation marker is more sensitive than the CD63 marker (C). DASA is 10-fold more sensitive than flow cytometry (D).

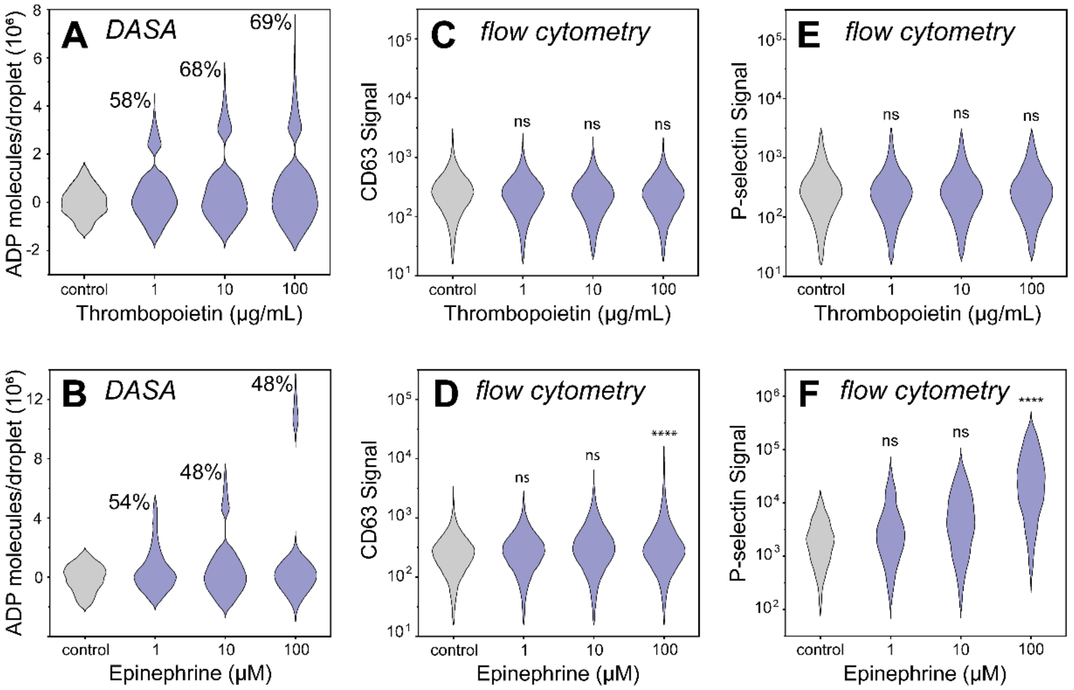


**SI Figure 4.** *Dose-dependent ADP secretion in response to priming.* DASA data of platelet responses to 1–100 ng/mL thrombopoietin (A) and 1–100 μM epinephrine (B). Flow cytometry using CD63 (dense granules) did not report responses to thrombopoietin (C) or epinephrine (D). Analysis using P-selectin (alpha granules) did not report responses to thrombopoietin (E) or 1–10 μM epinephrine (F). P-selectin only reported platelet activation (*P* <.0001) with the highest epinephrine concentration (100 μM). Significance relative to vehicle controls determined using a one-way ANOVA with Tukey's test.

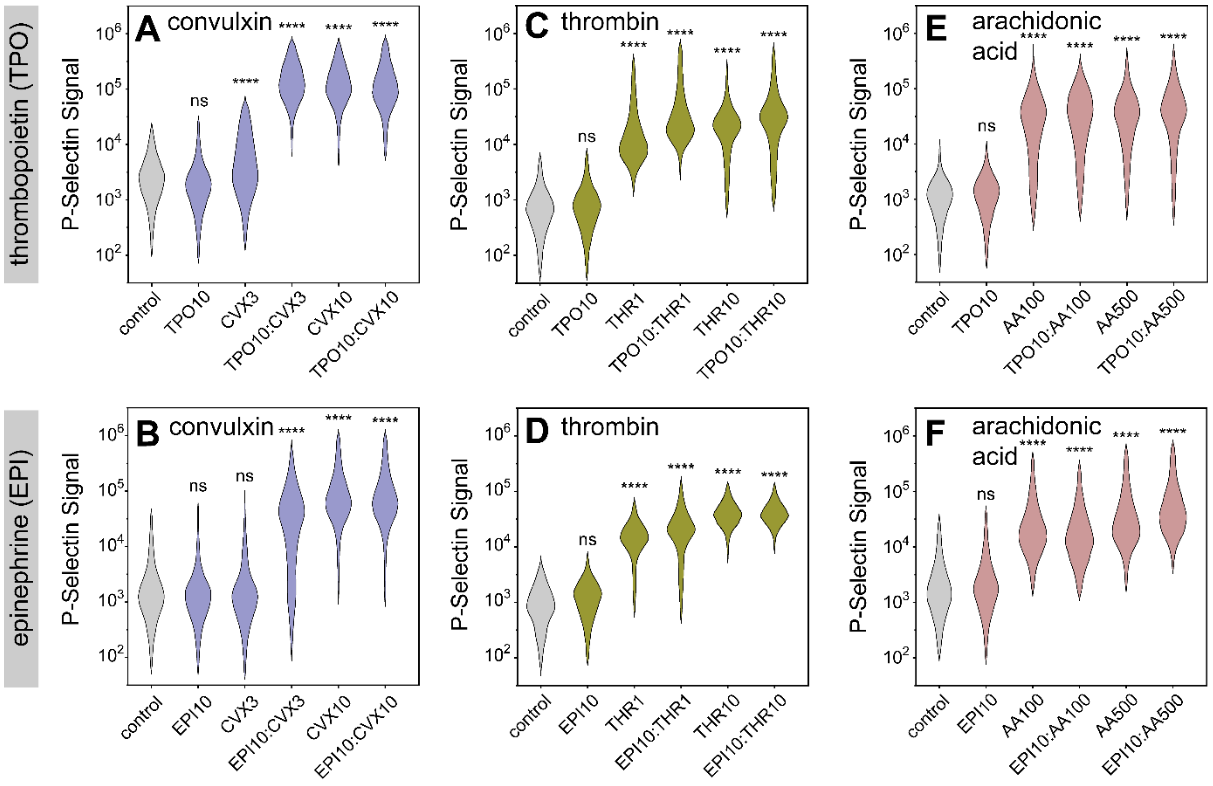


**SI Figure 5.** *Flow cytometry of platelets responses to a panel of agonists and primers.* P-selectin exposure was used as an activation marker, with platelets stimulated with convulxin (CVX, lilac, 3 or 10 ng/mL, A,B), thrombin (THR, green, 1 or 10 IU, C,D) and arachidonic acid (AA, pink 100 or 500 µM, E,F), and primed with either thrombopoietin (TPO, 10 µg/mL) or epinephrine (EPI, 10 µM). Significance relative to vehicle controls determined using a one-way ANOVA with Tukey's test.

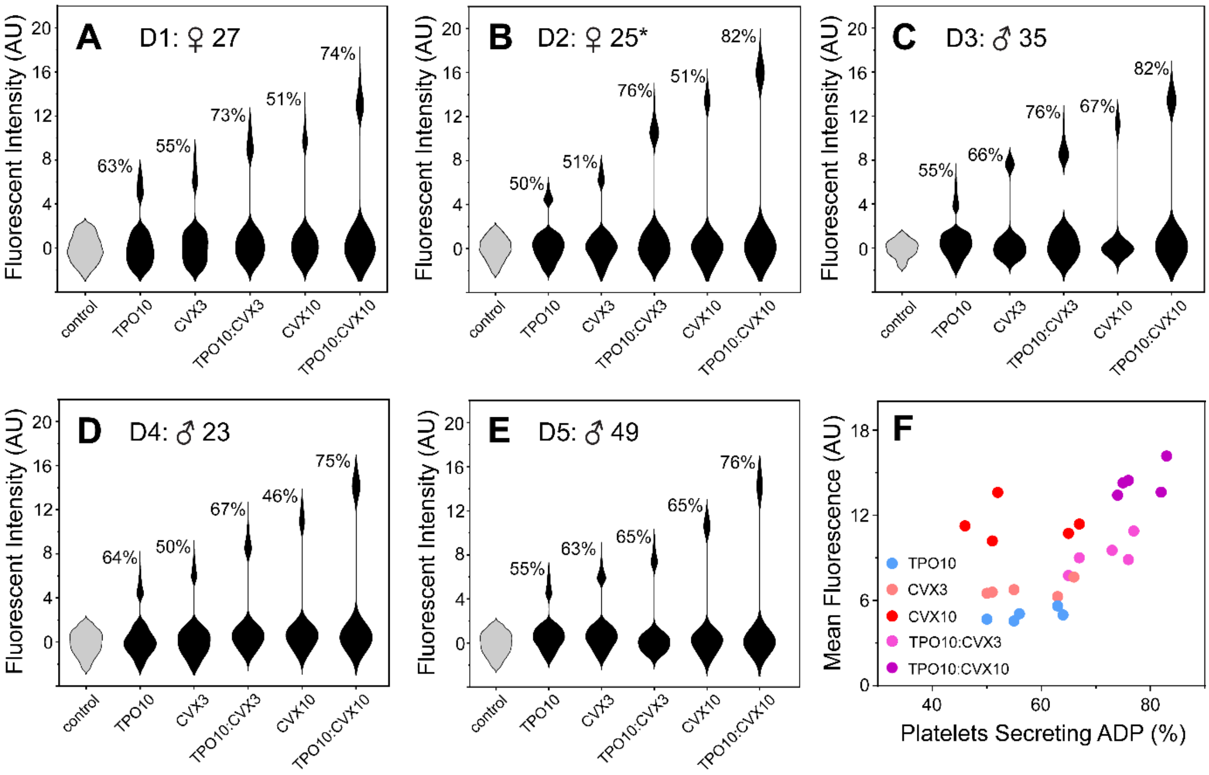


**SI Figure 6.** *Healthy donor variability.* ADP secretion levels and fraction of secreting platelets for different healthy donors (age, gender and contraceptive*, A–E). Platelets were treated with convulxin (CVX, 3 and 10 ng/mL), thrombopoietin (TPO, 10 μg/mL) and dual treatment. ADP secretion levels and fraction secreting platelets had low correlation (F).

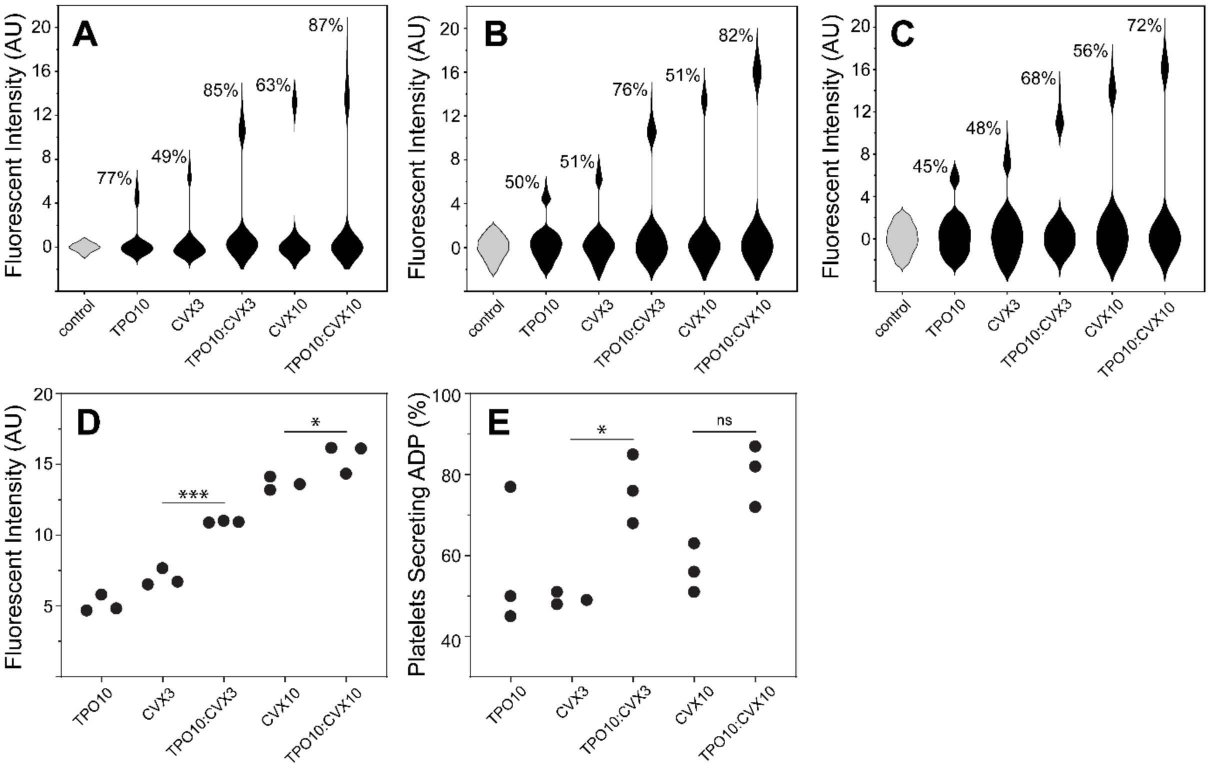


**SI Figure 7.** *Donor dynamics.* DASA monitoring of a healthy donor (25-year-old female) over a 4-month period (A–C). Platelets were treated with convulxin (CVX, 3 or 10 ng/mL) and thrombopoietin (TPO, 10 μg/mL). Consistent DASA responses to CVX and dual treatment with TPO and CVX were observed (D,E). Significance determined using a one-way ANOVA with Tukey's test.

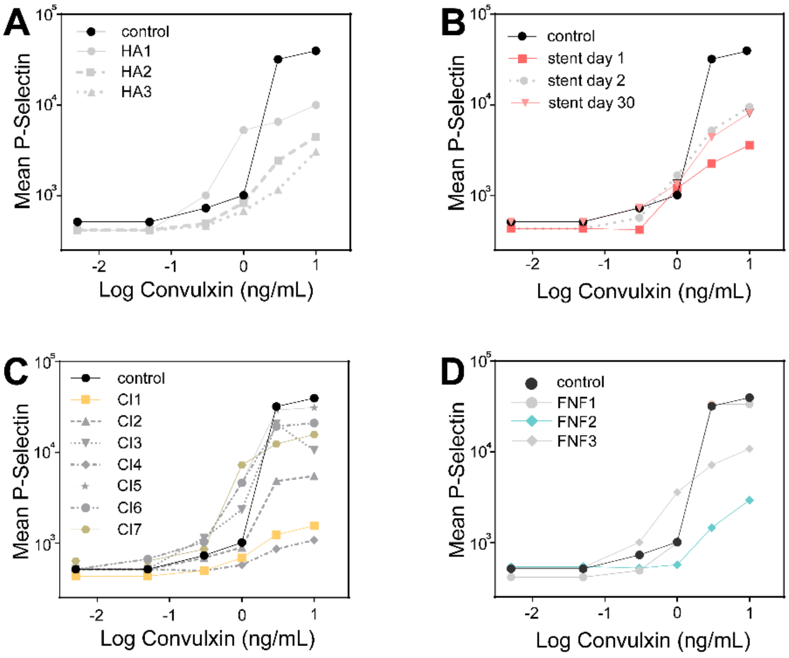


**SI Figure 8.** *Flow cytometry data from clinical cohorts.* Convulxin dose-response results using P-selectin as an activation marker. Acute myocardial infarction patients (A), a patient with acute myocardial infarction due to stent thrombosis monitored for 30 days (B), chest infection patients (C) and fractured neck of femur patients (D).

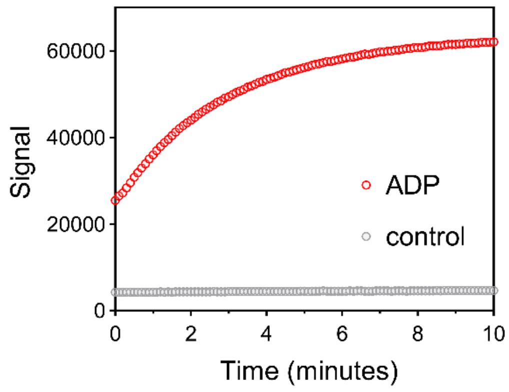


**SI Figure 9.** *ADP signal kinetics*. The fluorescent signal (Ex/Em; 535/587 nm) develops in 10 minutes.

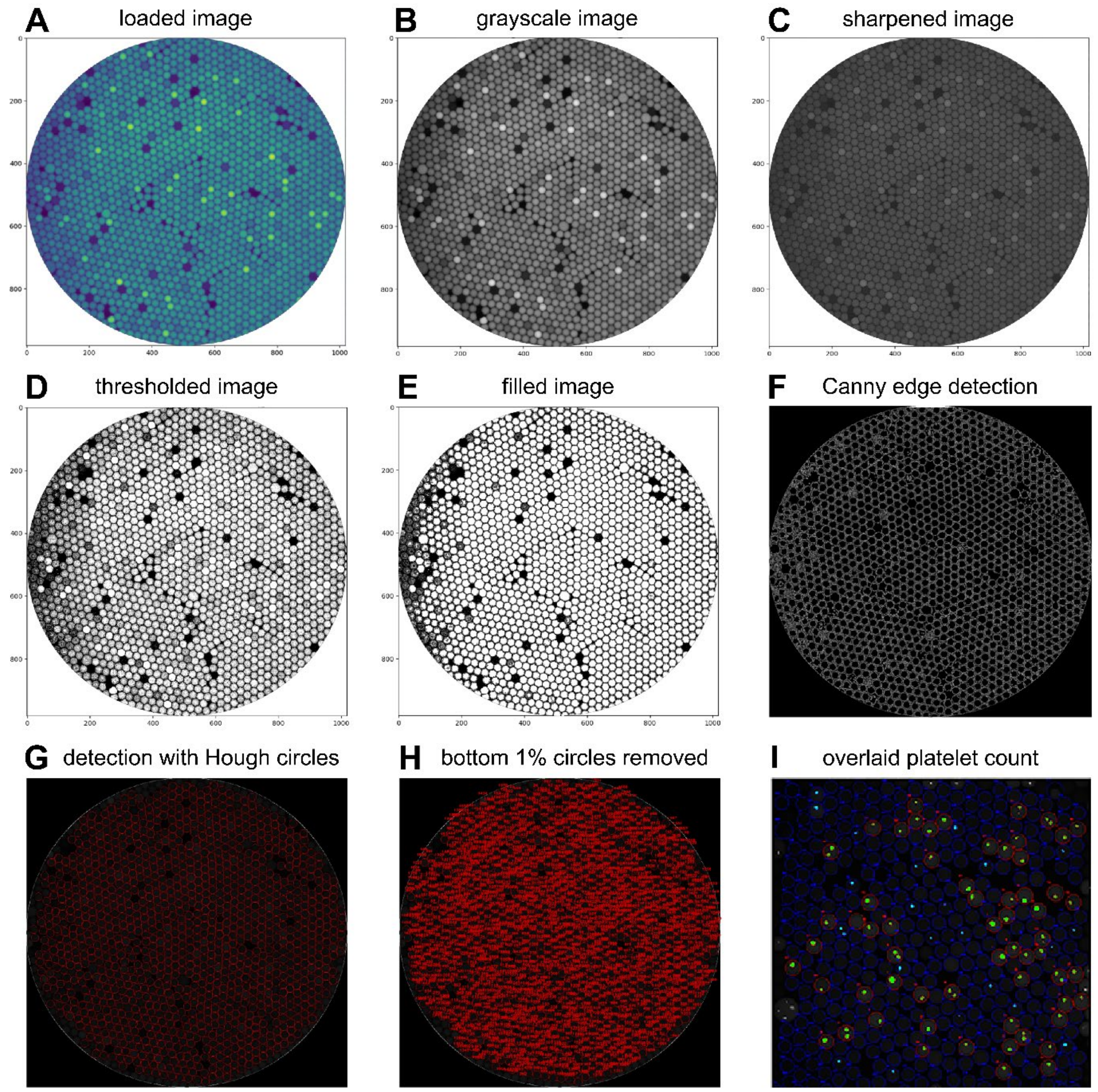


**SI Figure 10.** Illustrated Python image analysis pipeline to extract droplet fluorescent intensity and the platelet count.
